# Applying Lexical Link Analysis to Discover Insights from Public Information on COVID-19

**DOI:** 10.1101/2020.05.06.079798

**Authors:** Ying Zhao, Charles C. Zhou

**Affiliations:** Information Sciences Department, Naval Postgraduate School, Monterey, CA 93943, USA; Quantum Intelligence, Inc., Monterey, CA 93943, USA

**Keywords:** information mining, text mining, data mining, coronavirus, SARS-CoV-2, COVID-19, clade, mutation, genetics, tests, prevention

## Abstract

SARS-Cov-2, the deadly and novel virus, which has caused a worldwide pandemic and drastic loss of human lives and economic activities. An open data set called the COVID-19 Open Research Dataset or CORD-19 contains large set full text scientific literature on SARS-CoV-2. The Next Strain consists of a database of SARS-CoV-2 viral genomes from since 12/3/2019. We applied an unique information mining method named lexical link analysis (LLA) to answer the call to action and help the science community answer high-priority scientific questions related to SARS-CoV-2. We first text-mined the CORD-19. We also data-mined the next strain database. Finally, we linked two databases. The linked databases and information can be used to discover the insights and help the research community to address high-priority questions related to the SARS-CoV-2’s genetics, tests, and prevention.

**Significance Statement:** In this paper, we show how to apply an unique information mining method lexical link analysis (LLA) to link unstructured (CORD-19) and structured (Next Strain) data sets to relevant publications, integrate text and data mining into a single platform to discover the insights that can be visualized, and validated to answer the high-priority questions of genetics, incubation, treatment, symptoms, and prevention of COVID-19.

---

T his paper is to answer the call to action to the Nation’s data scientists and AI experts to mobilize and develop new text and data mining techniques that can help the science community answer high-priority scientific questions related to SARS-Cov-2, the deadly and novel virus, which has caused a worldwide pandemic and drastic loss of human lives and economic activities.

Recently, an open data set, namely the COVID-19 Open Research Dataset or CORD-19 (2) containing more than 40,000 full text scientific literature on SARS-CoV-2, was released (1, 2) for researchers across technology, academia, and the government. It is hopeful that machine learning and AI community might employ advanced data sciences to surface unique insights in the body of data and help answer with high-priority questions related to genetics, incubation, treatment, symptoms, and prevention. The corpus is updated weekly as new research is published in peer-reviewed publications and archival services like bioRxiv, medRxiv, and others. In this paper, we show how to apply an unique information mining method lexical link analysis (LLA) to link unstrurctured and structured data address the research questions below:

1. How to extract themes and topics that address the high-priority questions of genetics, incubation, treatment, symptoms, and prevention of COVID-19?
2. How to extract valued information such as information with authority, insights and innovation for combating SARS-Cov-2? What is the authoritative information and insightful information?
3. What are the timelines of themes and topics across all the research literature?

We show LLA that integrates text and data mining into a single platform so that insights from data can be visualized, and validated to answer the high-priority questions.

## Materials and Methods

### Overview of Lexical Link Analysis

Traditionally in social networks, the importance of a network node is a form of high-value information. For example, the leadership role in a social network (7, 8) is measured according to centrality measures (9). Among various centrality measures, a common practice is to sort and rank information based on authority. Current automated methods, such as graphbased ranking used in many search engines (10), require established hyperlinks, citation networks, social networks, or other forms of crowd-sourced collective intelligence. However, these methods are not applicable to situations where there exist no pre-established relationships among network nodes such as the data set SARS-Cov-2. This makes traditional methods difficult to apply. Furthermore, current methods mainly score popular information. High-value information can be totally different depending on specific applications. Popular and authoritative information identified by the current methods can be useful for marketing applications or crowdsourcing applications. Emerging and anomalous information is important for looking for insights and innovation. In paper, we show how to apply game-theoretic framework of lexical link analysis (LLA) to discover and rank high-value information from unstructured and structured data from SARS-Cov-2.

In LLA, a complex system can be expressed in a list of attributes or word features with specific vocabularies or lexicon terms to describe its characteristics. LLA is first a data-driven text analysis method. Fig. 1 shows an example of extracting and learning word pairs, or bi-grams as lexical terms, from text data. Words from a text document are represented as nodes, which form word pairs or bi-grams, via the links between any two nodes. For instance, the node “anitiviral” in Fig. 1 is formed with “chain-terminating,” “broadspectrum,” etc. as bi-gram word pairs (word features). In contrast to human-annotated word networks, such as WordNet (13), LLA automatically discovers word pairs, divides them into clusters and themes, and displays them as word networks. Fig. 1 shows an example of LLA.

**Fig. 1.**
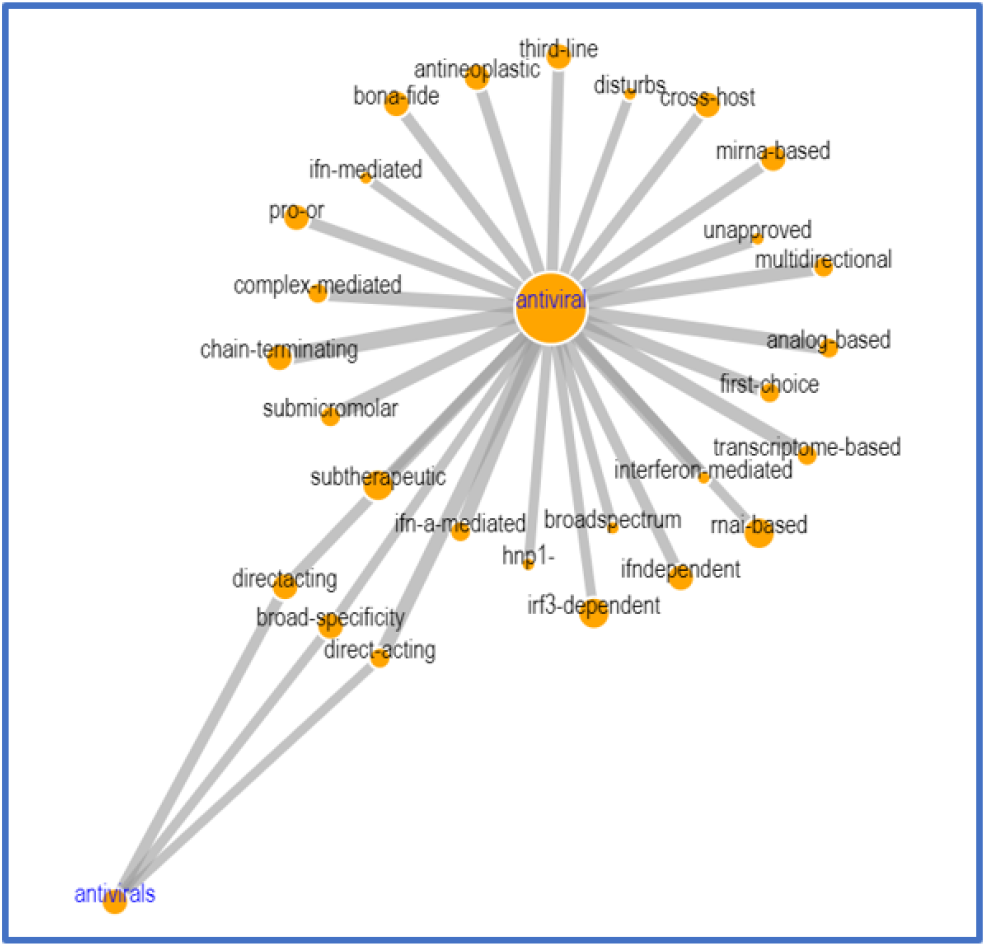
LLA Example

Bi-gram also allows LLA to be extended to structured data (15) including meta-data such as the ones for CORD-19, where a word is an attribute combined with its possible values. LLA is related to but significantly different from the methods such as bag-of-words (BOW) methods, Automap (7), Latent Dirichlet Allocation (LDA) (14), Latent Semantic Analysis (LSA) (16), Probabilistic Latent Semantic Analysis (PLSA) (17) and can be jointly used with NEE (20, 21), POS methods (25). LLA is related to unsupervised learning algorithms such as k-means, principal component analysis (PCA), and spectral clustering (18). In a social network, the most connected nodes are typically considered the most important nodes. However, the uniqueness of LLA is that we consider anomalous information (word features) might be more interesting. Community detection algorithms have been illustrated by Newman (11, 12) in terms of a quality function as the “modularity” measure for a community (cluster) and optimized using a dendrogram-like greedy algorithm. The uniqueness of LLA includes new value metrics to identify high-value information that are not presented in the other methods.The new value metrics consider a game-theoretic framework (19, 35).

### LLA Applied to CORD-19

LLA automatically discovers popular (P) themes, emerging (E) themes, and anomalous (A) themes (as defined below) as follows:

- Popular (P) themes: These themes resemble themes generated from the eigenvalue centrality measure in the network sciences. The themes represent the main topics in a data set. They can be insightful information in two folds: 1) These word pairs are more likely to be shared or cross-validated across multiple diversified domains, so they are considered authoritative; 2) These themes could be less interesting because they are already in public consensus and awareness and can be considered as popular.
- Emerging (E) themes: These themes tend to become popular or authoritative over time. An emerging theme has the intermediate number of inter-connected word pairs. They are emerging and important themes and can be high-value for further investigation.
- Anomalous (A) themes: These themes may not seem to belong to the data domain when compared to other themes. They are interesting outliers and can be high-value for further investigation.

Fig. 2 shows an example of extracted themes from a text data set where popular themes, e.g. 517(P), emerging theme, e.g. 42(E), and anomalous theme e.g. 478(A) among others are listed. Fig. 8 shows a drill-down visualization for theme 517(P) labeled “infection coronavirus, global coronavirus” The labeled nodes in Fig. 8 are the words with the most connections with other words (via bi-gram measures in LLA).

**Fig. 2.**
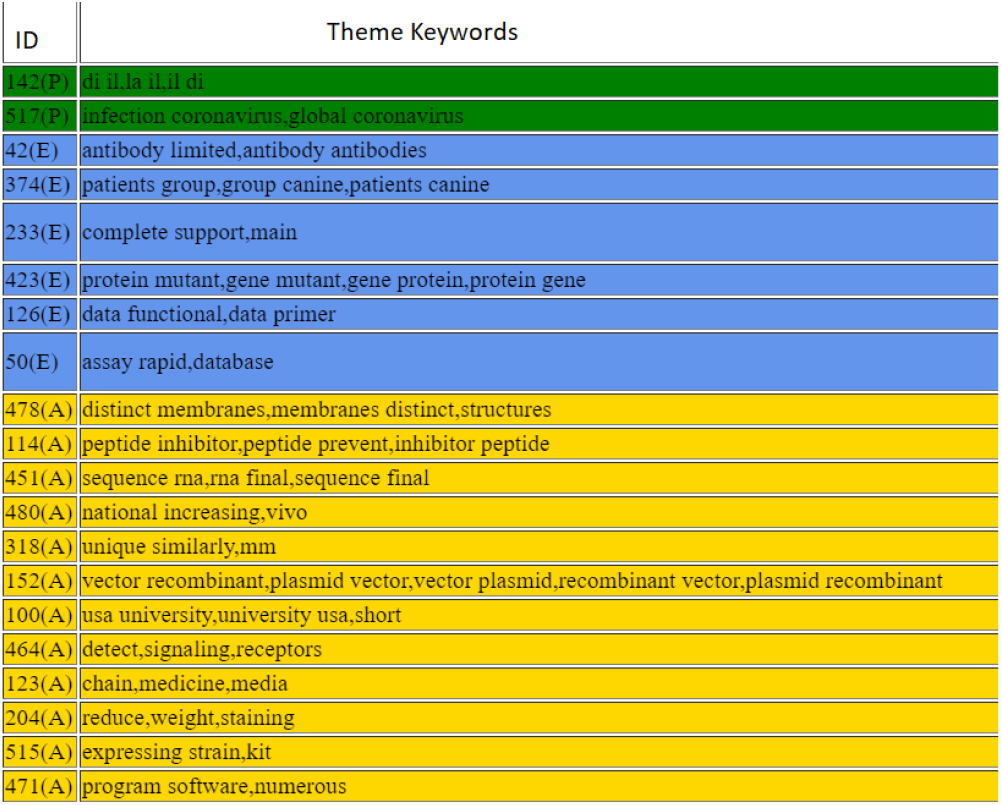
LLA themes

## The Next Strain Database

### Data

Understanding the spread and evolution of a virus such as SARS-Cov-2 is important for effective public health measures and surveillance. A global strain tracking tool from nextstrain.org (4, 6), i.e., which strain evolves from which other strain. Nextstrain consists of a database of viral genomes, a bioinformatics pipeline for phylodynamics analysis of many novel virues such as Zika, Ebola, and SARS-Cov-2 with an interactive visualization platform. The visualization integrates sequence data with other data types such as geographic information, serology, or host species. Nextstrain compiles the current understanding of phylogenetic analysis into a single accessible location, open to health professionals, epidemiologists, virologists and the public alike (5).

### Insights

Fig. 3 shows the phylogenetic tree of the SARS-Cov-2’s similarity and development around the globe from 12/3/2019 to 3/25/2020 where the 2649 cases of patients’ genomic data uploaded to the nextstrain.org website. Visually, Fig. 3 shows the strain in North America and Europe mutated to a different branch of the tree around 2/25/2020 and is more contagious. Fig. 4 shows the same data set colored by clades. Tab. 1 shows a split of the 2649 cases to three time periods: before 2/25/2020, from 2/25/2020 to 3/16/2020, after 3/16/2020. We computed the average case per day for Clade A2, A2a (the new mutation) and Clade A1a and other types. Clade A2, A2a is the mutated strain because it only has the data starting from 1/28/2020. Clade A1a, etc. and A2, A2a are statistically different (*p* < 0.0001) measured via average cases per day for each time period, i.e., 6.29 vs 0.24 (before 2/25/2020), 38.81 vs 46.86 (between 2/25/2020 and 3/16/2020), and 17.89 vs 31.33 (after 3/16/2020). A2a almost started on 2/25/2020 since only 11 cases of A2, A2a happened before 2/25/2020. Clade A2,A2a is more contagious than Clade A1a, and other since the average cases per day is much higher, 46.86 and 31.33 between 2/25/2020 and 3/16/2020. After 3/17/2020, many countries in North America and Europe for both clades, however, the mutated clade decreased statistically significantly slower (31.33) than A1a, B1, B2, etc. together (17.89).

**Fig. 3.**
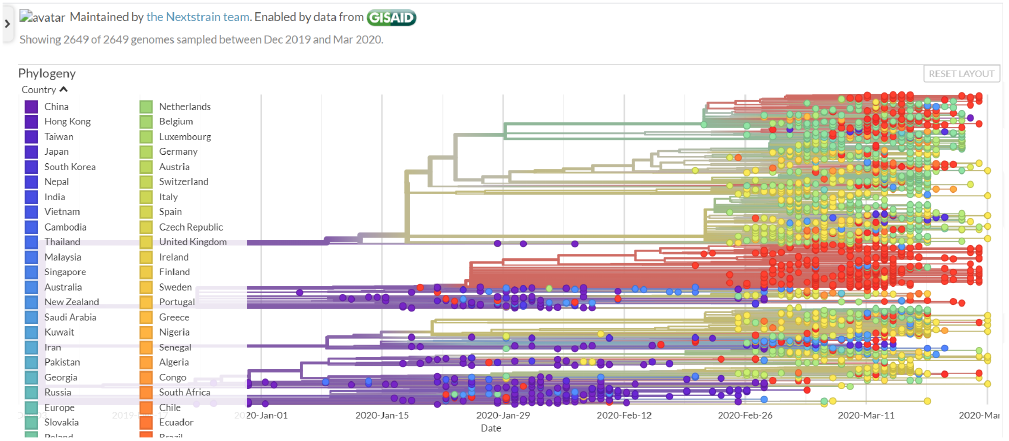
Next strain data by country from 12/3/2019 to 3/25/2020. Source: nextstrain.org

Fig. 5 shows how the first case of Clade A2a linked to earlier cases of Clade A2. There are two insights as follows:

- Among the total 1277 cases of Clade A2, A2a, there were only 34 cases in Asia, including four cases in China and 18 cases in Japan. There were 955 cases in Europe, 216 cases in North America, and 15 cases in Australia.
- Among the four cases of Clade A2, A2a in China, three were collected from 1/28/2020 to 2/6/2020, they were submitted around 3/20/2020. The fourth case was collected on 3/23/2020 and submitted at the same time.

**Fig. 4.**
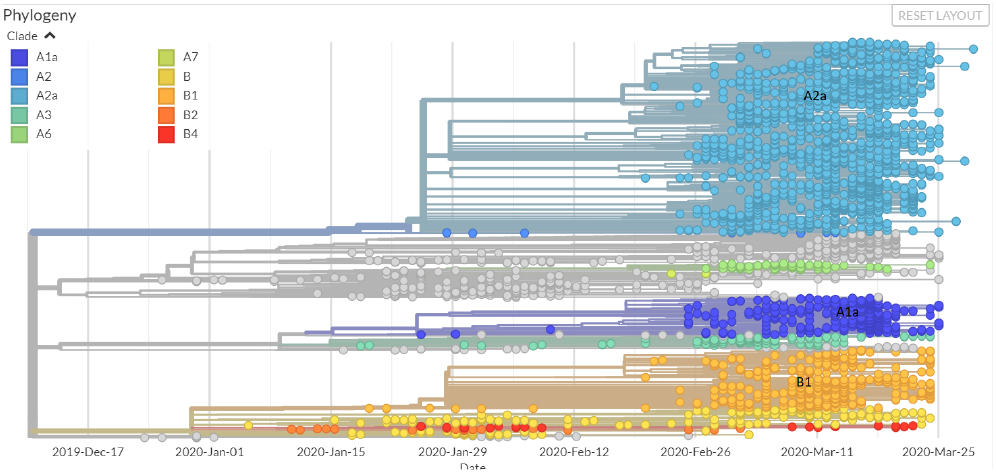
Next strain data by clade from 12/3/2019 to 3/25/2020. Source: nextstrain.org

**Fig. 5.**
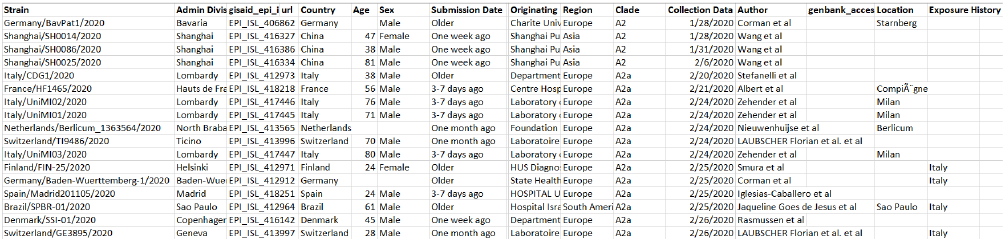
The first case of Clade A2a. Source: nextstrain.org

**Table 1.** Cases

| A1a, B1, B2, etc | Cases | Days | Cases/Per Day |
| --- | --- | --- | --- |
| 12/24/2019-2/24/2020 | 396 | 63 | 6.29 |
| 2/25/2020-3/16/202 | 815 | 21 | 38.81 |
| 3/17/2020-3/25/2020 | 161 | 9 | 17.89 |
| A2, A2a |  |  |  |
| 1/28/2020-2/24/2020 | 11 | 46 | 0.24 |
| 2/25/2020-3/16/202 | 984 | 21 | 46.86 |
| 3/17/2020-3/25/2020 | 282 | 9 | 31.33 |

At least the data shows the Clade A2, A2a in Europe and North America are more contagious and virulent.

### Linking the Next Strain Data Set to the CORD-19 Data Set

The next strain data can be linked to the CORD-19 data set using LLA. We first extracted another document data set with documents pertinent to the next strain database. We then fused the two data sources based on themes in Fig. 2.

Fig. 6 shows a match matrix of number of matched word pairs from two document sources (the next strain and CORD-19), there are 2012 matched word pairs. Fig. 7 shows the examples of matched word pairs. These matched word pairs (concepts) are also grouped based on these themes. Fig. 8 shows a popular theme of word pair appeared in both data sources. Fig. 9, Fig. 10, and Fig. 11 show examples of emerging themes appeared in both data sources. Fig. 12 shows an example of an anomalous theme appeared in both data sources. Emerging themes are interesting topics for researchers to drill down and discover information in CORD-19 pertinent to high-priority questions of the SARS-Cov-2 of genetics (Group 423(E)), tests (Group 50(E)), and prevention (Group 42(E)).

**Fig. 6.**
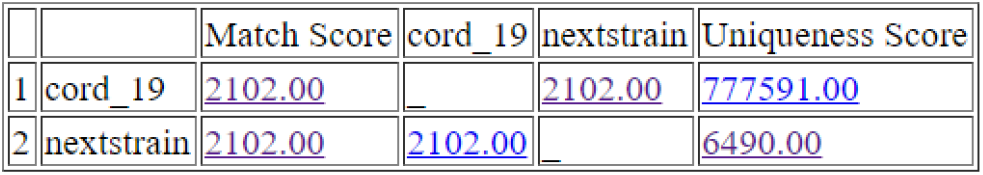
LLA match matrix

**Fig. 7.**
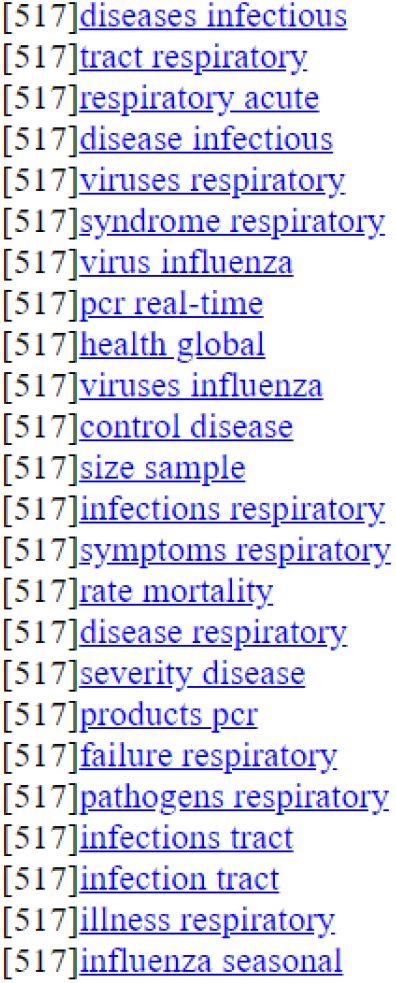
LLA matched list

**Fig. 8.**
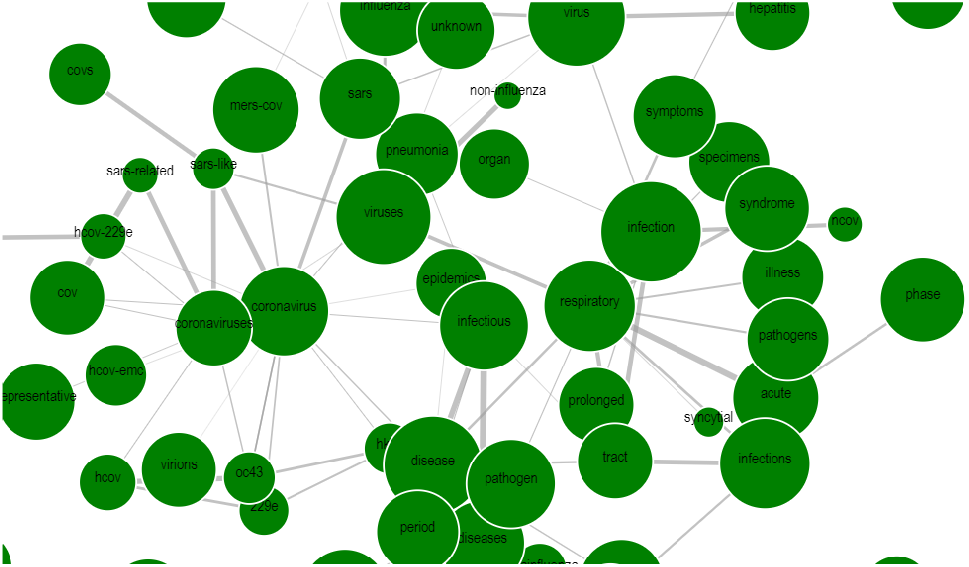
LLA group 517(P)

**Fig. 9.**
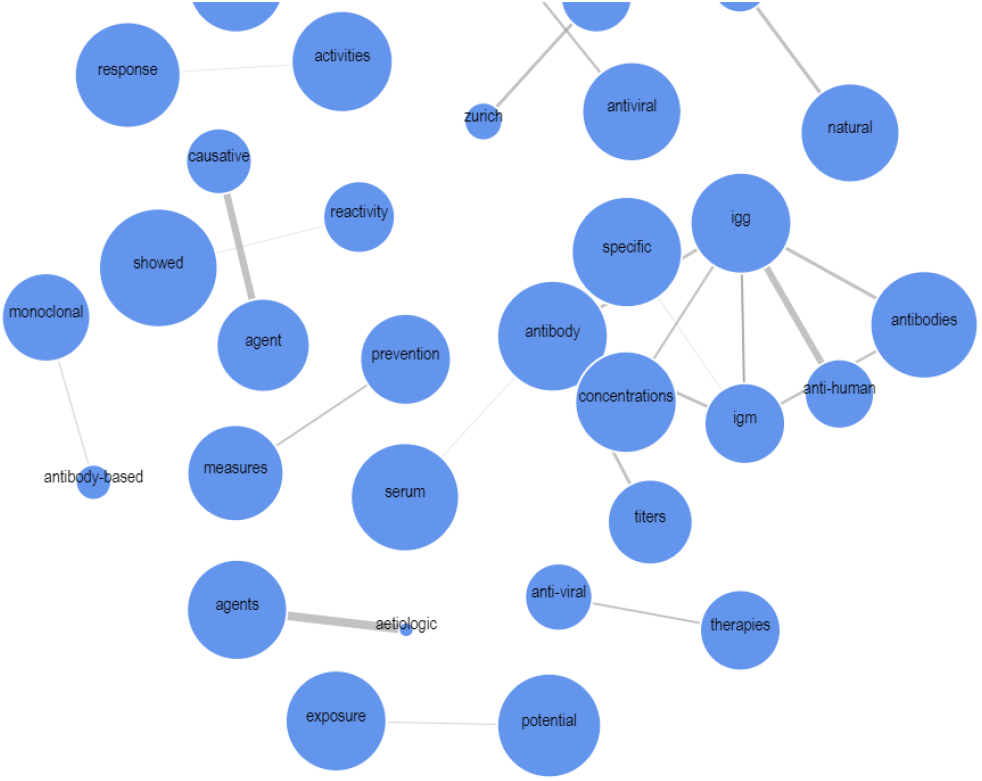
LLA group 42(E)

**Fig. 10.**
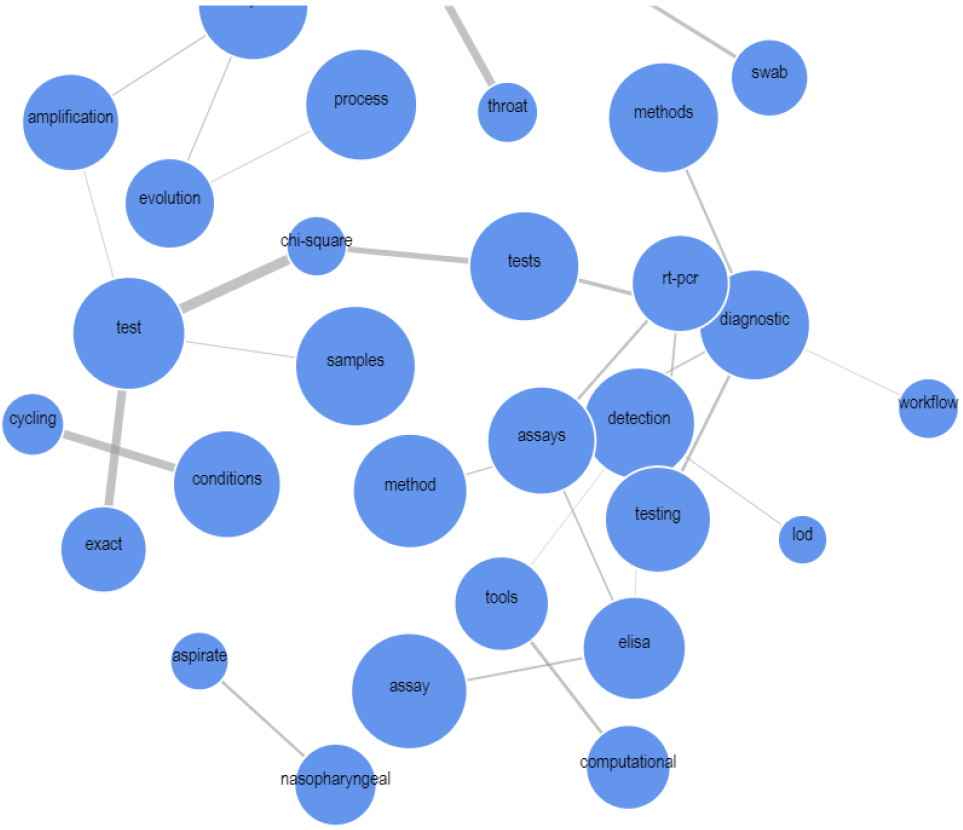
LLA group 50(E)

**Fig. 11.**
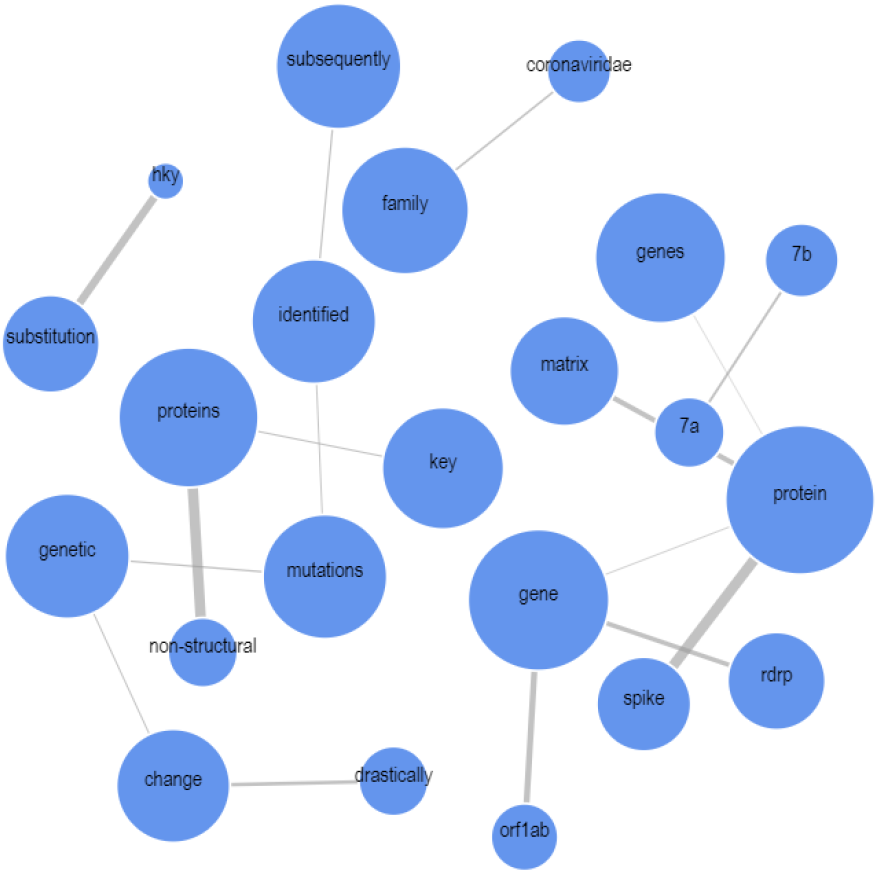
LLA group 423(E)

**Fig. 12.**
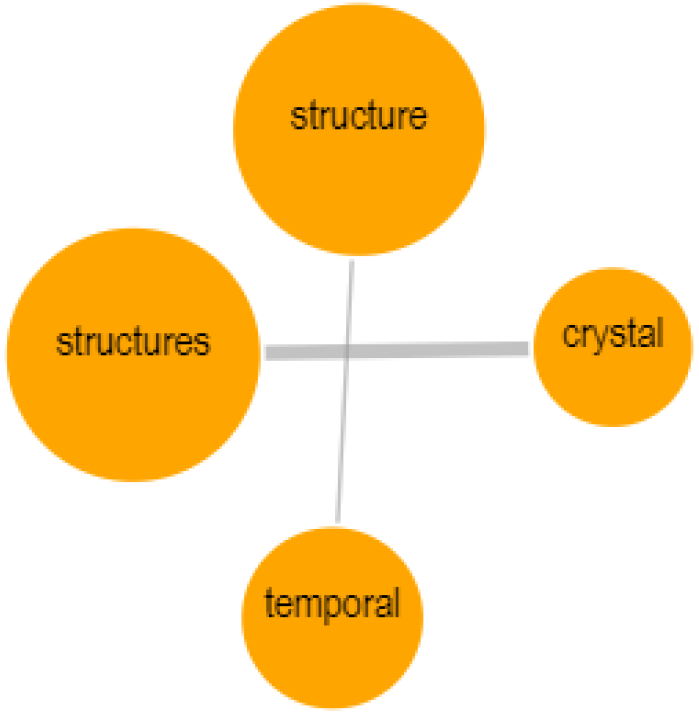
LLA group 478(A)

### Conclusion and Challenge to Future

We applied an unique information mining method lexical link analysis to conduct a preliminary study to the call to action and help the science community answer high-priority scientific questions related to SARS-Cov-2. We first text-mined an unstructured database, i.e. the COVID-19 Open Research Dataset or CORD-19. We also data-mined a structured database, i.e., the next strain database, covering the period from 12/3/2019 to 3/25/2020. Finally, we linked two databases and certain publications, and discovered the insights and methodologies. The linked documents from CORD-19 can help address high-priority questions related to SARS-COV-2’s genetics, tests, and prevention.

There are some un-answered questions we need to ponder:

1. If certain data and publications before 12/3/2019 need to be mined and analysed, can one estimate the outcome of COVID-19 outbreak after 12/3/2019?
2. Can we use the data and publications up to today to estimate what will happen one month later?

